# SFCalculator: connecting deep generative models and crystallography

**DOI:** 10.1101/2025.01.12.632630

**Authors:** Minhuan Li, Kevin Dalton, Doeke Hekstra

## Abstract

Proteins drive biochemical transformations by transitioning through distinct conformational states. Understanding these states is essential for modulating protein function. Although X-ray crystallography has enabled revolutionary advances in protein structure prediction by machine learning, this connection was made at the level of atomic models, not the underlying data. This lack of connection to crystallographic data limits the potential for further advances in both the accuracy of protein structure prediction and the application of machine learning to experimental structure determination. Here, we present SFCalculator, a differentiable pipeline that generates crystallographic observables from atomistic molecular structures with bulk solvent correction, bridging crystallographic data and neural network-based molecular modeling. We validate SFCalculator against conventional methods and demonstrate its utility by establishing three important proof-of-concept applications. First, SFCalculator enables accurate placement of molecular models relative to crystal lattices (known as phasing). Second, SFCalculator enables the search of the latent space of generative models for conformations that fit crystallographic data and are, therefore, also implicitly constrained by the information encoded by the model. Finally, SFCalculator enables the use of crystallographic data during training of generative models, enabling these models to generate an ensemble of conformations consistent with crystallographic data. SFCalculator, therefore, enables a new generation of analytical paradigms integrating crystallographic data and machine learning.

## 1 Introduction

Machine learning (ML) is making rapid strides in the prediction of biomolecular structures and their conformational ensembles. Key capabilities include the ability to predict structure from sequence with near-experimental accuracy[4, 30, 36], to design practically viable structures from geometric or functional constraints[14, 19, 66], and to predict ensembles of possible conformations given an initial structure[29, 35]. The capability to generate credible conformations as a function of a sequence, sequence alignment, or a latent variable overcomes some sampling limitations inherent in stepwise Molecular Dynamics and Monte Carlo algorithms.

X-ray crystallography, on the other hand, has enabled experimental structure determination for many biomolecules, including assemblies such as ribosomes, nucleosomes, and proteasomes, and a wealth of smaller systems such as enzymes, ion channels, GPCRs, and kinases. Spurred on by powerful new X-ray Free Electron Lasers[49, 61] and synchrotron beamlines[26, 58], X-ray crystallography has also begun to enable visualization of the dynamics of proteins on timescales from femtoseconds to seconds—for example visualizing enzyme catalysis[13, 60], photosynthesis[6], K^+^ ion channel permeation[34], and the first events in vision[24]—and across physical and chemical conditions—for example enabling crystallographic screening of interactions of drug targets with thousands of drug fragments[62].

Generative ML models and X-ray crystallographic data could be productively combined to improve the accuracy of generative models, and the accuracy and throughput of crystallographic structure determination. Recent work has taken important steps in this direction. McCoy et al.[40] showed that AlphaFold models are often sufficiently accurate to provide an initial solution to the crystallographic phase problem. Terwilliger et al.[64] showed that by iteratively feeding intermediate structure refinement models as templates into the AlphaFold module one can achieve higher-quality models. This iterative procedure is necessary, however, because there was no direct interface between predictive models and crystallographic data.

Differentiable algorithms that integrate cryo-electron microscopy (cryo-EM) and nuclear magnetic resonance (NMR) data with machine learning have significantly accelerated structure determination, boosting both efficiency and throughput[33, 69]. Here, we introduce SFCalculator, a fast, differentiable likelihood-calculation tool that connects structural models—complete with solvent corrections—to crystallographic observables. SFCalculator is designed as a bridge between the rapidly evolving machine learning ecosystem and crystallographic datasets, as illustrated in Fig. 1.

**Fig. 1.**
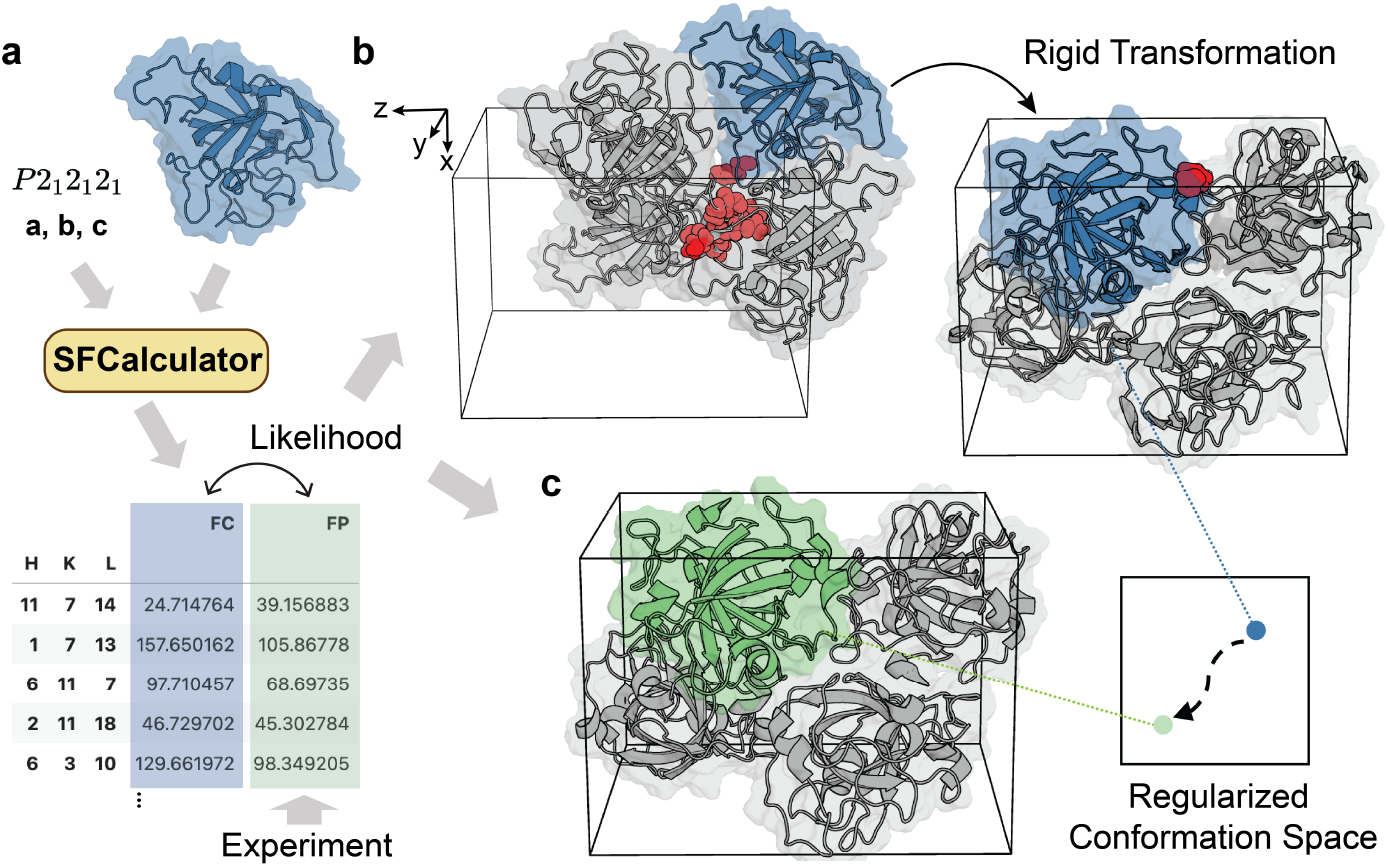
SFCalculator as an interface. SFCalculator provides a differentiable forward function that calculates from molecular models, along with space group and unit cell information, expected values of experimental observables. This enables the computation of a differentiable likelihood or scoring function (**a**), facilitating downstream tasks such as molecular replacement (**b**), and structure refinement (**c**) within a unified framework. The red spheres in panel (**b**) represent atoms that clash with neighboring atoms.

We validate SFCalculator by benchmarking it against conventional calculations and demonstrate its utility through three examples. First, we show how SFCalculator can be combined with a hierarchical grid search algorithm[68, 69] and gradient-based optimization [32] to place a search model (here, an AlphaFold prediction, Figure 1a) relative to a protein crystal lattice, a procedure known as molecular replacement (Figure 1b). We then asked if the latent space of a pre-trained conformational generative model, the Boltzmann Generator [47], could serve as a search space to find a protein conformation maximally consistent with experimental data (Figure 1c). By doing so, we find that AlphaFold models can be refined to yield conformations with excellent stereochemistry and fit to the data.

Finally, we show how SFCalculator can be used to train generative models on crystallographic data, offering, at once, a perspective on the further improvement of generative models, and a principled strategy for tackling the crystallography inverse problem: inference of conformational ensembles that exhibit both strong data fit and favorable physical energies. These approaches naturally incorporate constraints encoded in the generative model design, potentially obviating the need for explicit physical restraints[25, 44] during refinement.

## 2 Results

### SFCalculator provides a fast and differentiable interface

SFCalculator relates atomic models of protein structure to the primary crystallographic observables. Specifically, X-ray scattering methods measure the amplitudes of the Fourier components (structure factors) of the electron density of a sample. SFCalculator, likewise, provides access to the Fourier components of the electrostatic potential map and thus, in principle, supports electron scattering methods like cryo-electron microscopy and electron diffraction—we will address this further in future work.

The electron density of a protein crystal includes contributions from both macro-molecular atoms and the surrounding solvent. The disordered solvent component is particularly significant in low-resolution data. A widely implemented approach for modeling this solvent contribution is the bulk-solvent model, which presumes a uniform solvent density in all regions not occupied by the atomic structure. A more refined variant, known as the probe-shrink model[28], permits variations in solvent density within a boundary region at the interface between the solvent and macromolecule to account for the ordered hydration layer. This model has become the default in contemporary crystallographic refinement programs[3, 45].

Despite its utility, the discretization and rounding operations inherent in this approach hinder differentiability, thereby precluding the simultaneous optimization of the macromolecular and bulk-solvent components during refinement. To address this, Fenn et al.[18] introduced a differentiable algorithm that smooths the solvent boundary using Gaussian or polynomial functions. However, this method is computationally intensitve and relies on Babinet’s principle[43], neglecting the influence of the hydration layer. In SFCalculator, we have designed algorithms to ensure that both contributions are incorporated in a fast and differentiable manner, as illustrated in Fig.2 (a). We adopt the direct summation method for the macromolecular contribution and a cutoff-sigmoid approximation of the solvent mask (see Methods). To assess the accuracy and performance of SFCalculator, we first performed a rigorous comparison with the widely used software PHENIX [3]. For a comprehensive evaluation, the comparison was conducted across a large sample of PDB entries (*N* =868), covering data resolutions from 0.8 to 3.0 Å, various spacegroups, and a range of molecular sizes (see Fig. A1). The results demonstrate strong agreement with PHENIX across key metrics, while showcasing the computational efficiency and scalability of SFCalculator.

A comparative analysis of solvent masks was performed between the probe-shrink method[28], the default in PHENIX, and the differentiable solvent mask generated by SFCalculator. Fig. 2(b) presents slice views of the probe-shrink mask, as comparison to our differentiable mask slice in Fig. 2(a). While the overall shapes of the two masks closely agree, notable differences are evident in the boundary regions. The probe-shrink method provides more nuanced details in these areas, primarily due to its “shrink” step, which excludes the hydration layer surrounding the macromolecule. This adjustment is crucial, as solvent molecules in the hydration layer are often ordered and are not adequately represented by a flat solvent model.

**Fig. 2.**
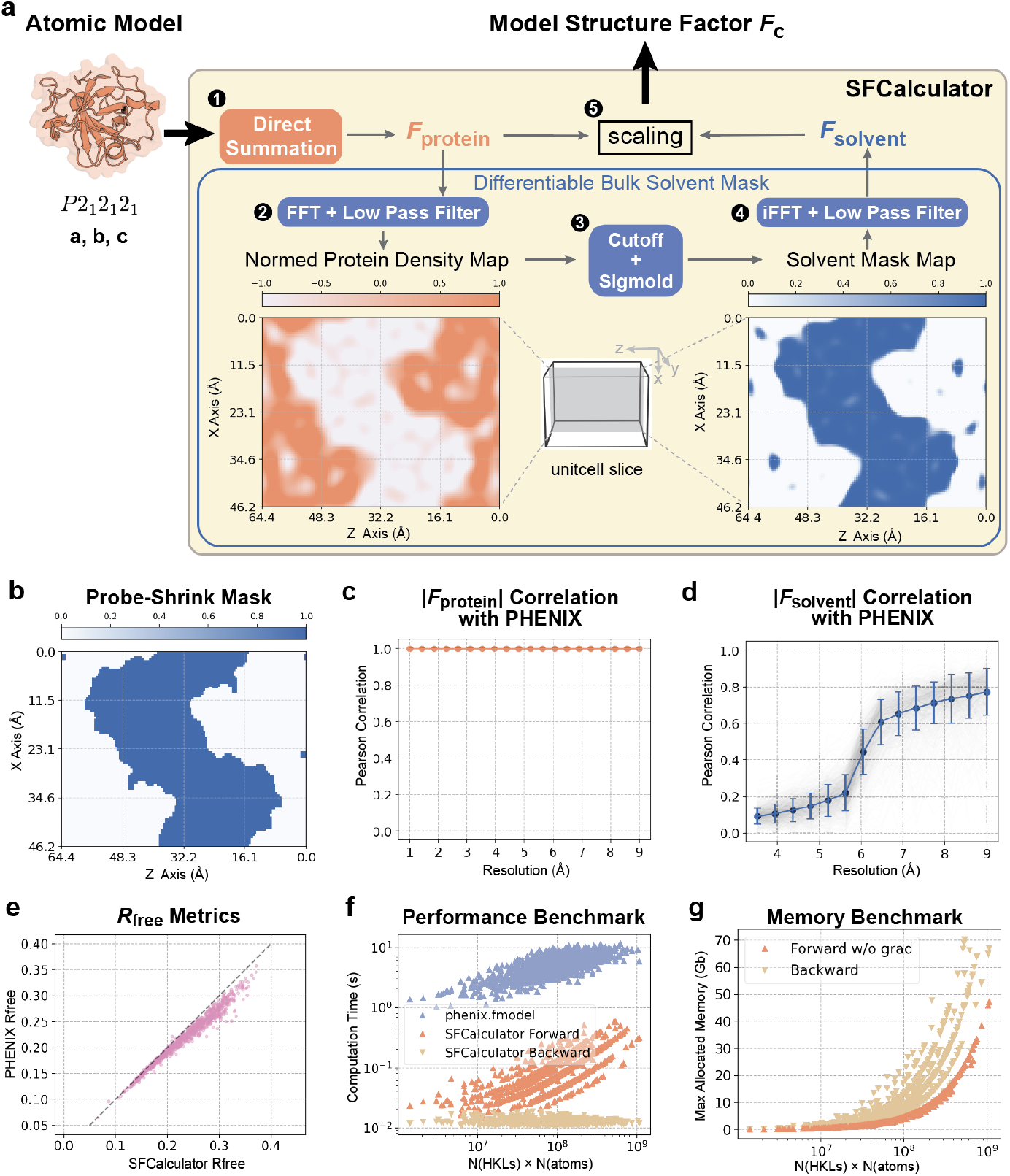
Algorithms and benchmark of SFCalculator. (**a**) Algorithms of SFCalculator: The protein contribution is computed through direct summation, generating **F**_protein_. The differentiable bulk solvent mask algorithm is then applied to compute the solvent contribution, **F**_solvent_. These contributions are combined by determining parameterized scale factors to produce the final model structure factor *F*_*c*_. (**b-g**) Accuracy and performance benchmarking of SFCalculator compared to the commonly used software PHENIX. (**b**) Slice view of the solvent mask generated using the Probe-Shrink method (default in PHENIX) compared to our differentiable solvent mask in (**a**). (**c**) Correlation statistics of protein structure factors between SFCalculator and PHENIX, showing perfect agreement. (**d**) Correlation statistics of solvent structure factors between SFCalculator and PHENIX. (**e**) Comparison of *R*_*free*_ calculations between SFCalculator and PHENIX. SFCalculator achieves high consistency overall, with discrepancies appearing for high-*R*_*free*_ datasets. (**f**) Performance benchmarking of SFCalculator’s forward and backward computations compared to PHENIX’s phenix.fmodel. SFCalculator is 50-200 times faster due to GPU utilization, while PHENIX runs solely on CPUs. Hardware details: PHENIX was tested on an Intel(R) Xeon(R) CPU @ 2.20 GHz, and SFCalculator was tested on an Nvidia A100 GPU. (**g**) Memory benchmarking of SFCalculator, demonstrating the ability to handle proteins with up to ∼1,000 residues at -2 Å resolution (PDB id: 4PKF) on a single Nvidia A100 80GB GPU.

To assess the agreement between structure factors generated by SFCalculator and PHENIX, we calculated their correlation. For the protein component, SFCalculator shows perfect correlation with PHENIX (Fig. 2c), demonstrating the accuracy of its calculations. For the solvent component (Fig. 2d), high correlations are observed at low resolutions (high Å values) with less correlation at high resolutions due to differences in handling of the protein-solvent boundary.

We further compared the quality of structure factors calculated for the same deposited models by SFCalculator and PHENIX, using the cross-validation metric *R*_free_ to relate structure factors calculated by both to the data (see Methods). Overall, SFCalculator achieves high consistency, with discrepancies primarily for dataset/-model combinations with high *R*_free_ (Fig. 2e). These discrepancies are primarily due to the larger contribution of solvent at low resolution, where differences between the solvent models of SFCalculator and PHENIX become more pronounced. Despite these differences, SFCalculator maintains robust performance across a wide range of resolutions.

SFCalculator is computationally efficient. In performance tests comparing forward and backward computations, SFCalculator achieved speeds 50–200 times faster than PHENIX’s phenix.fmodel (Fig. 2f), which operates exclusively on CPUs. This large performance gain derives from the ability of GPUs to leverage parallel processing and the differentiable backends integrated into SFCalculator.

We further evaluated the memory efficiency of SFCalculator (Fig. 2g). We find that SFCalculator can handle proteins with approximately 1000 residues at a resolution of 2 Å (PDB id: 4PKF) on a single Nvidia A100 80GB GPU. Handling even larger systems can be achieved by further partitioning certain bottleneck operations that involve large matrices with *N*_*HKL*_ *× N*_*atoms*_ elements. This scalability underscores the potential for large-scale applications in structural biology, making SFCalculator a valuable tool for analyzing complex macromolecular systems.

Taken together, these results establish the accuracy, efficiency, and scalability of SFCalculator, positioning it as a promising complement to established tools like PHENIX, for workflows requiring high throughput and differentiability.

### SFCalculator Enables Efficient Molecular Replacement

As a first test of SFCalculator’s potential to facilitate crystallographic data processing, we used it to implement a simple molecular replacement (MR) algorithm. MR is used in crystallography to find an initial, approximate solution to the phase problem and is practically useful when the target molecule shares structural similarity with a previously solved model. With the advent of highly successful protein structure prediction algorithms, many targets that were previously challenging or intractable– —particularly those involving novel folds—–can now be readily solved using MR [5, 31, 40, 42]. The immediate goal of MR is to establish the position and orientation of a protein relative to the crystal lattice—inherently, a 6-dimensional search. MR has been extensively studied, resulting in the development of several efficient programs[23, 39, 41] based on a decomposition of the 6D search into subsequent rotational and translational searches, comparing observed and calculated Patterson functions as proposed by Rossmann and Blow [56]. Our implementation follows the same principles but incorporates a hierarchical rotational grid constructed using Hopf Fibration [68, 69] (see Methods).

We illustrate molecular replacement using SFCalculator using a crystallographic dataset of the inverting cellulase PcCel45A, a fungal endoglucanase (PDB id: 3×2I) with a “Newton’s cradle proton relay mechanism in its active site[46]. This enzyme crystallizes in an orthorhombic space group, P 2_1_2_1_2_1_, with symmetry-related hydrolase molecules within the unit cell. Each molecule represents a distinct ground-truth solutions for the pose search (Fig. 3 a).

**Fig. 3.**
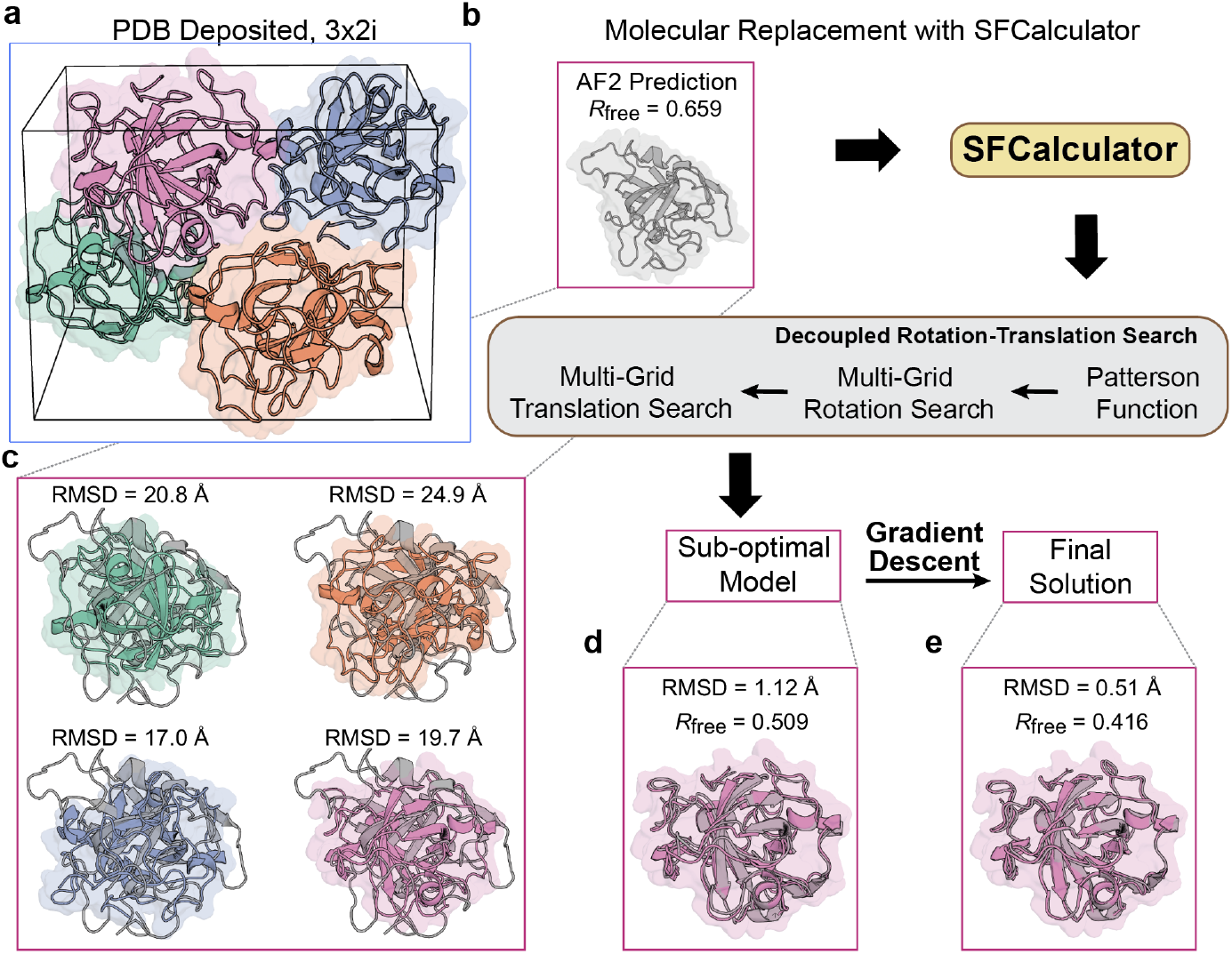
Molecular Replacement using SFCalculator. (a) Visualization of the PcCel45A unit cell (PDB id: 3×2I), which contains four copies of the asymmetric unit, providing four ground-truth poses for the pose search. (b) Visualization of the AlphaFold-predicted structure for the enzyme system. The random initial pose assigned by the predictive model yields a poor *R*_free_ value (0.659), indicating a significant mismatch. (c) The initial pose exhibits a large R.M.S.D. (Root-Mean-Square Deviation) relative to each of the four ground-truth poses, with values ranging from 17.0 Å to 24.9 Å. (d) The hierarchical grid search identifies an approximate pose, closely resembling the pink ground-truth pose, with an improved R.M.S.D. of 1.12 Å and an *R*_free_ of 0.509. (**e**) Subsequent gradient descent refinement further improves the R.M.S.D. to 0.51 Å and reduces *R*_free_ to 0.416, underscoring the importance of the gradient descent step in achieving accurate pose refinement.

The molecular replacement procedure begins with an initial model generated by AlphaFold2. Due to the random pose assignment of the predicted model, the starting R_free_ is extremely poor (0.659), as shown in Fig. 3 b, as is the Root-Mean-Square Deviation (R.M.S.D.) between the initial pose and any of the four ground-truth poses, ranging from 17.0 to 24.9 Å (Fig. 3c).

To find an approximate pose, we then employed a hierarchical grid search in decoupled rotation and translation spaces. This search identified a poses similar to one of the ground-truth poses (specifically, the pink copy), resulting in an R.M.S.D. of 1.12 Å and *R*_free_ of 0.509 (Fig. 3d). Subsequent gradient descent, relying on autodifferentiation enabled by SFCalculator, strongly improves R.M.S.D., to 0.51Å, and *R*_free_, to 0.416, as shown in Fig. 3e. It is remarkable that, even after multiple rounds of hierarchical grid search—–with the final round featuring a 0.95 degree step for rotation and 0.75 Å step for translation, gradient descent can still achieve such large improvements just by improving pose. This observation highlights a key limitation of hierarchical grid search: sensitivity to the initial candidates. If the initial base grid is not sufficiently fine, the search is prone to converging on local minima. As the computational cost of increased grid resolution scales cubically in step size, finer grids are impractical, underscoring the value of the final gradient descent step.

### SFCalculator Enables Structure Refinement Using Generative Models

Initial structural models of a protein for an experimental condition of interest are often fraught with inaccuracies–—errors that can obscure critical details of molecular mechanisms. Structure refinement addresses this by optimizing structural parameters such as atomic positions and atomic displacement factors (e.g., B factors). Structure refinement using experimental data generally faces two fundamental obstacles. First, experimental data rarely suffice to yield physically accurate models without the use of additional stereochemical information on, e.g. bond lengths, angles, and preferred side chain orientations, during model building. Second, the energy landscapes of proteins are extremely high-dimensional and ‘rugged’, with many local minima separated by barriers.

Current approaches to structure refinement typically address the need to include stereochemical information by combining a data term with explicit restraint terms in the overall objective function to be minimized. These restraints can be derived from high-resolution protein and small-molecule crystal structures[25], quantum calculations[38], or molecular mechanics forcefields [27][9][44]. Efficient navigation of conformational landscapes is typically achieved through a heuristic combination of different sampling and gradient-descent steps using several model representations [3].

We reasoned that conformational generative models could help address both major obstacles by enabling alternative approaches to protein structure refinement. Generative models are often trained to transform coordinates from internal “latent-space” representations with relatively simple properties (e.g., following a Gaussian distribution) to complex real-world representations (e.g., protein atomic structures). As such, their latent spaces might be more readily traversed during structure refinement than traditional coordinate representation spaces, while their coordinate transforms may have learned to account for protein physics, enabling optimization of model fit to experimental data (a likelihood function) without the use of explicit constraints or restraints on conformations.

SFCalculator makes it possible to test this idea by providing a fully (auto)differentiable link between models and experimental likelihood evaluations. To do so, we adopted the normalizing flow architecture[15, 50] as the foundational framework for our model, building on the Boltzmann Generator approach pioneered by Noé and colleagues[47]. In this approach, we learn a transformation *x* = *T*_θ_(*z*), from latent variable z to protein coordinates *x. z* follows a smooth, easily sampled distribution, such as a Gaussian (*z ∼ µ*(*z*) = 𝒩 (0, *I*)). Under the change of variables formula, the induced distribution in *x*-space can then be expressed as

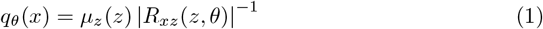

where *R*_*zx*_ represents the determinant of the Jacobian matrix of the transformation *T*_θ_(*z*). The objective is to learn model parameters θ such that the variational distribution *q*_θ_(*x*) approximates the true Boltzmann distribution of protein conformations. A key advantage of the normalizing flow framework is its inherently invertible transformation design, which ensures compatibility with any initial structure model, allowing seamless integration without introducing deviations. The schematic architecture is composed of several key blocks, which we describe with detail in Appendix B.4.

We reimplemented the Boltzmann Generator of Noé et al., as depicted in Fig. 4a, employing two complementary training objectives based on correspondence with molecular dynamics (MD) snapshots and physical energy calculations for self-generated snapshots, as described in Methods. We illustrate the results with the same PcCel45A dataset used to illustrate Molecular Replacement. As intended, the generated samples displayed reasonable energies (Fig. 4c). We then froze the parameters θ of the Boltzmann Generator.

**Fig. 4.**
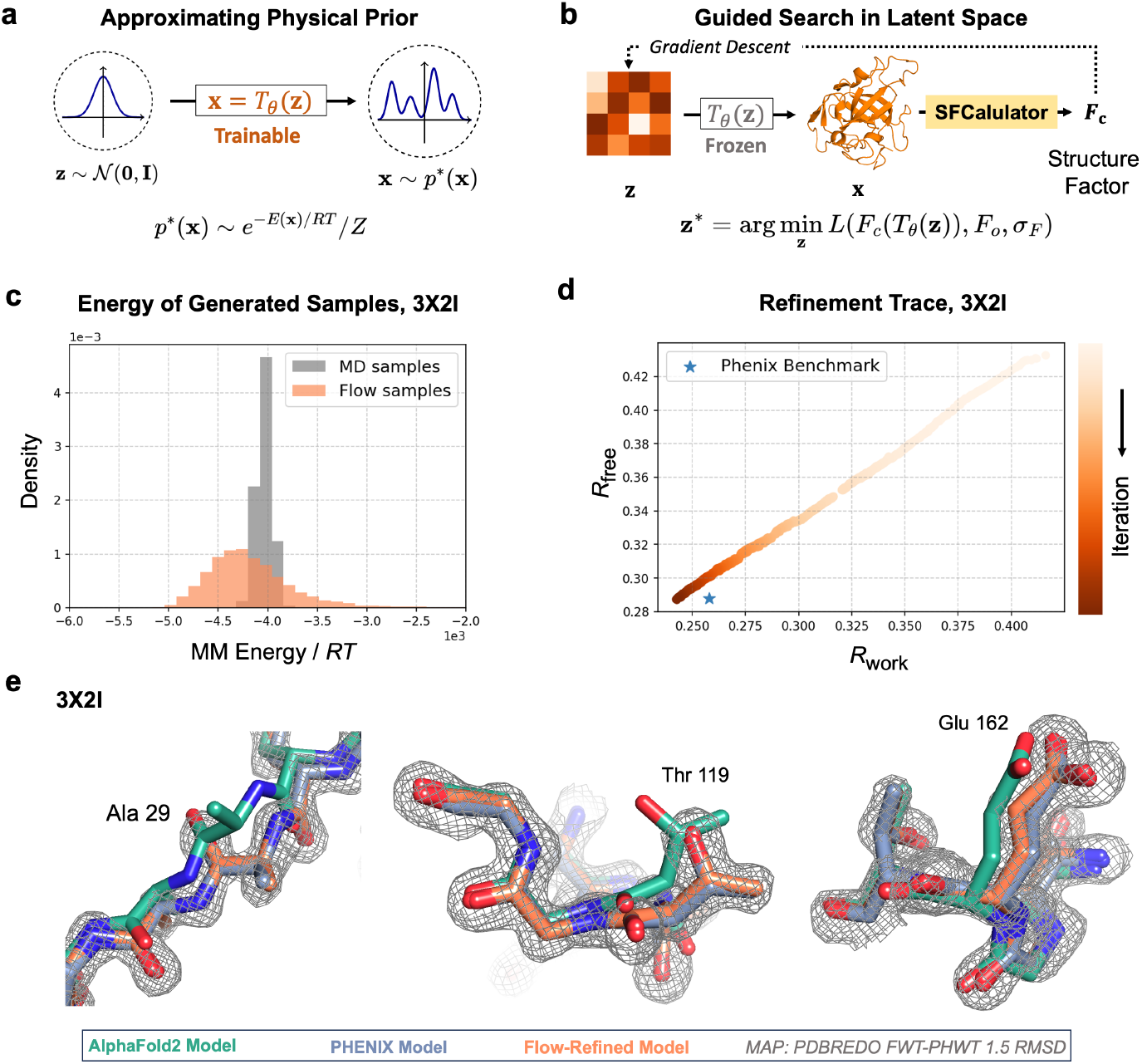
Guided search in the latent space of a generative model encoding a physical prior. (**a**) The physical prior, represented by the Boltzmann distribution *p*^∗^(*x*) = *e*^−*E*(*x*)*/RT*^ */Z* at temperature *T* for conformations *x* with energy *E*(*x*), is approximated using a normalizing flow model. This framework embeds the latent space *z* with the physical prior, where *z* ∼ 𝒩 (0, *I*), and the transformation *x* = *T*_*θ*_(*z*) maps the latent space to the target space *x* to approximate the target distribution *p*^∗^(*x*). (**b**) After training, the refinement process is performed through guided search in the latent space. Gradient descent is applied to minimize the data likelihood loss, *L* (*F*_*c*_ (*T*_*θ*_(*z*)), *F*_*o*_, *σ*_*F*_), using SFCalculator to compute structure factors (*F*_*c*_). (**c-e**) Refinement of the PcCel45A structure (PDB id: 3×2I) in a Latent Space with Embedded Physical Priors. (**c**) Energy distribution of samples generated by the trained normalizing flow, compared to samples from molecular dynamics (MD). The generated samples exhibit energies within a reasonable scale, consistent with physical expectations. Refinement trace in the latent space. (**e**) Visualization of the improvements achieved through refinement in the latent space. The method effectively corrects both backbone and side chain conformations, showcasing structural improvements over the initial models.

Next, we examined whether the latent space of this generative model would allow for efficient refinement of protein structure by unconstrained stochastic gradient optimization of the experimental data likelihood (see Methods) given the model, as illustrated in Fig. 4b. Importantly, the differentiability of SFCalculator enables gradient information to flow back, guiding the search in the latent space **z**.

The optimization trajectory is shown in Fig. 4d. Our method achieved a data fit, measured by *R*_free_, similar to the model refined by PHENIX, while the molecular mechanics energy from our refinement (*−*4, 105 *kT*) was significantly more reasonable than that of the PHENIX-refined model (+6, 584 *kT*). This difference primarily arises from PHENIX’s use of a limited set of geometric restraints [25]. While effective, these restraints do not accurately account for features such as atomic clashes, and hydrogen bonding and electrostatic interactions.

In Fig 4e, we visualize the structural improvements achieved through our refinement process, on both the protein backbone and side chains. These results validate the utility of SFCalculator and demonstrate the potential of using deep generative models for structural refinement.

### SFCalculator Enables Ensemble Refinement by Generative Modeling

In the above example, we first trained a generative model and then used its latent space as a search space to find a conformation that best accounted for the experimental data. SFCalculator also makes it possible to directly include crystallographic data during training of generative models. Notably, of course, many conformational generative models, including AlphaFold [30], have been trained on atomic models derived from crystallographic data, but none have been trained on the crystallographic data themselves. As a first example of doing so, we include crystallographic data during the training of a Boltzmann Generator in order to obtain conformational ensembles consistent with crystallographic data. Training a generative model this way serves as a form of ensemble refinement[10], yielding a variational approximation [55] of the true posterior distribution of conformations of a protein consistent with physical prior information and the crystallographic data.

As illustrated in Fig. 5a, the target distribution follows the unnormalized posterior distribution:

**Fig. 5.**
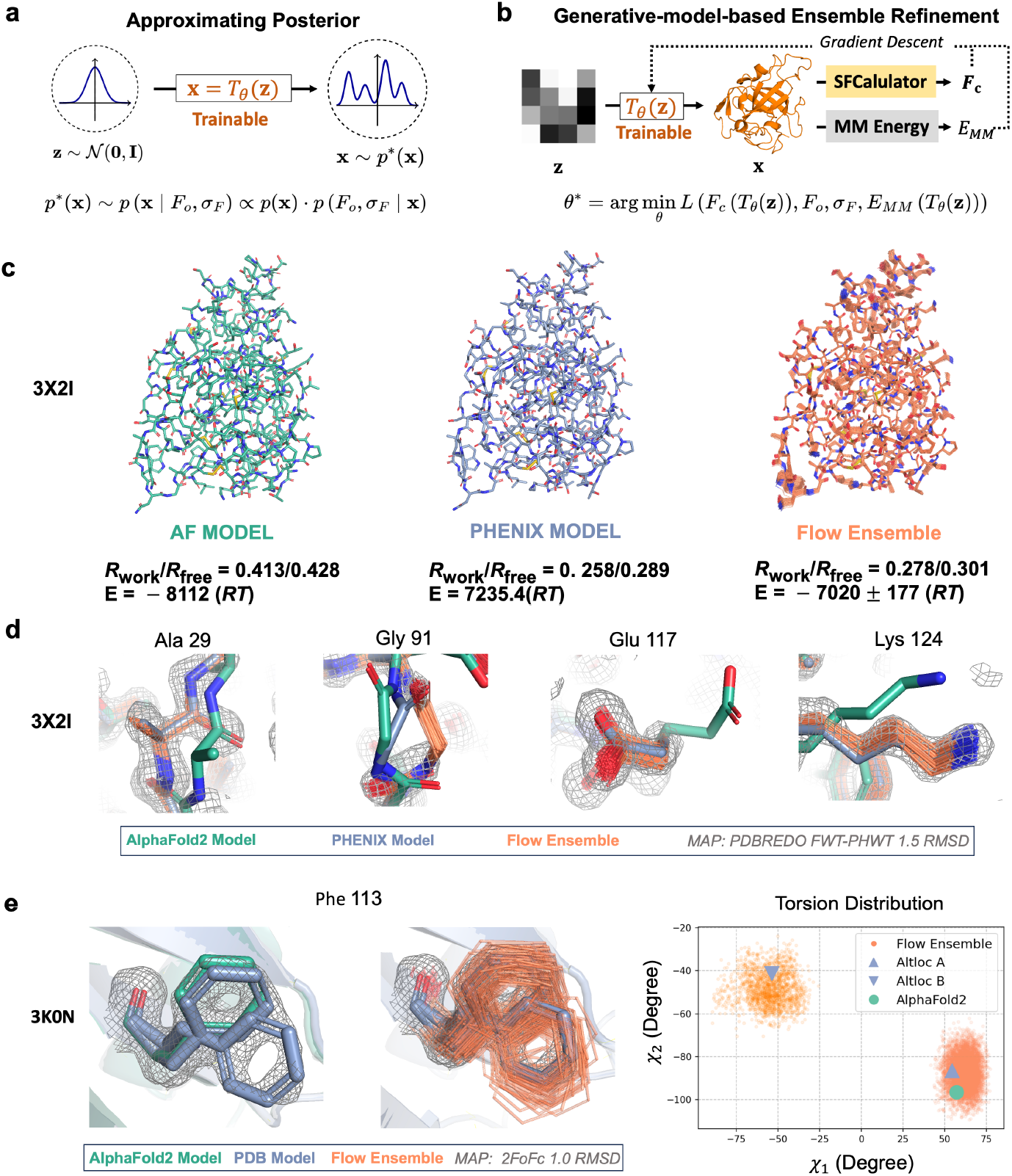
Ensemble refinement using a generative model approximating the posterior distribution. (a) The target distribution of the normalizing flow is the posterior combining the physical prior *p*(*x*) and the experimental likelihood. The transformation *x* = *T*_*θ*_(*z*) is trained to approximate this posterior. (**b**) By including both the experimental likelihood and the molecular mechanics (MM) energy in the training objective, the parameters *θ* are optimized via gradient descent. This ensures that the samples from the normalizing flow align with the posterior distribution. SFCalculator computes structure factors (*F*_*c*_), while the MM energy term (*E*_*MM*_) incorporates physical constraints, refining the ensemble to balance data fit and physical plausibility. (**c**) Comparison of models for PcCel45A (PDB ID: 3×2I). The AlphaFold2 model after molecular replacement exhibits a high *R*_work_/*R*_free_ = (0.413 / 0.428) but a good molecular mechanics (MM) energy -8112 RT. The Phenix-refined model achieves better *R*_work_/*R*_free_ =(0.258 / 0.289) but has an unphysically high MM energy 7235.4 RT. The flow ensemble model strikes a balance with *R*_work_/*R*_free_ =(0.278 / 0.301) and a more reasonable MM energy E=-7020 ± 177 RT. (**d**) Visualization of key residues (ALA29, GLY91, GLU117, and LYS124) of PcCel45A. The AlphaFold2, Phenix, and flow ensemble models are overlaid with electron density maps, showcasing the structural improvements achieved by the flow ensemble refinement. Both backbone and side chain conformations are corrected, providing better agreement with the experimental data. (**e**) Refinement results for residue PHE113 in the isomerase Cyclophilin A (CypA, PDB ID: 3K0N). The 3*k*0*n* dataset, collected at room temperature, is known to exhibit alternative conformations for many residues, including PHE113. The flow ensemble successfully captured both conformational states. The torsion angle distribution plot for *χ*_1_ and *χ*_2_ highlights the broader conformational diversity achieved by the flow ensemble.

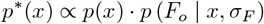

where *p*(*x*) represents the physical prior Boltzmann distribution, and *p* (*F*_*o*,_ | *x*, σ_*F*_) corresponds to the experimental likelihood of the observed structure factor amplitudes *F*_*o*_ given the protein coordinates *x*, and measurement uncertainty σ_F_. Once the normalizing flow is trained, the model itself serves as an approximation of the posterior distribution. Consequently, samples generated from the model can be interpreted as effective results of ensemble refinement.

We evaluated this ensemble refinement approach on two systems: the hydrolase PcCel45A used above, and the proline isomerase Cyclophilin A (CypA, PDB ID: 3K0N). The results demonstrate the advantages of approximating the posterior distribution with flow-based generative models to achieve physically meaningful and experimentally consistent conformational ensembles.

For the hydrolase, as shown in Fig. 5c, the flow ensemble models strikes a balance between consistency with the data (*R*_work_/*R*_free_ =0.278 / 0.301) and a more reasonable MM energy E = -7,020 *±* 177 *kT*. Further analysis of key residues highlights the structural improvements achieved by flow-based ensemble refinement, as illustrated by Fig. 5d for residues ALA29, GLY91, GLU117, and LYS124. When overlaid with electron density maps, the models refined by the flow ensemble show corrected backbone and side chain conformations, providing good agreement with the experimental data.

We further tested the refinement method on Cyclophilin A (CypA, PDB ID: 3K0N), a system known for exhibiting an important network of alternative conformations of residues along its central beta sheet [20], including PHE113. The flow ensemble successfully captures both conformational states of PHE113, as shown by the overlay with the electron density map in Fig. 5e. Additionally, the torsion angle distribution plot for χ_1_ and χ_2_ of PHE113 reveals the broader conformational diversity achieved by the flow ensemble compared to other models. This demonstrates the ability of the flow ensemble to capture diverse and physically meaningful conformational states.

## 3 Discussion

A defining strength of SFCalculator is its differentiability, which seamlessly connects crystallographic data with machine learning models for protein structure generation. This capability enables gradient-based optimization, allowing crystallographic observables to be integrated into end-to-end machine learning pipelines for joint refinement, structural validation, and iterative model improvement. Its scalability ensures compatibility with large datasets and high-throughput workflows, positioning SFCalculator as a critical bridge between traditional crystallographic methods and emerging computational approaches. This integration accelerates advancements in protein structure prediction and generation, which are addressing long-standing challenges in structural biology and opening new avenues for understanding molecular function and interactions.

Despite advances in modern refinement algorithms, significant challenges remain, particularly in automation. The rugged, non-convex optimization landscape of macro-molecular refinement often leads algorithms to local minima. Overcoming this requires careful initialization and iterative strategies, such as simulated annealing, to navigate toward globally optimal solutions. However, automated methods still struggle to replicate the nuanced, context-dependent decisions made during expert manual inspection. Tasks like identifying alternative conformations or resolving ambiguities in poorly defined regions often rely on visual intuition and expertise[11, 17]. This reliance on manual intervention creates bottlenecks, limiting scalability for high-throughput datasets and underscoring the need for more adaptive and automated algorithms.

The complexity of the optimization landscape is closely tied to its representation. Transformations between coordinate systems, such as Cartesian and dihedral angles, can affect landscape navigability without eliminating inherent barriers. Generative modeling approaches in machine learning are designed to identify and exploit such transformations[7, 15, 47], facilitating the principled incorporation of diverse prior constraints. This makes the integration of generative models into refinement workflows particularly promising.

Leveraging the differentiability of SFCalculator, we demonstrate proof-of-concept applications that enhance the interpretation of crystallographic data within generative models of protein structure. By incorporating physical priors from molecular mechanics force fields–—rather than relying solely on geometric constraints derived from small-molecule statistics—–we establish a more physically grounded refinement framework. The combination of SFCalculator with deep generative models also enables rigorous variational inference, producing conformational ensembles that not only fit the experimental data but also maintain favorable physical energies. This synergy underscores the potential of integrating machine learning with crystallographic refinement to drive progress in structural biology.

### Refinement of atomic displacement parameters

While the present work focuses on conformational refinement, two critical aspects of structural refinement remain unaddressed: the refinement of atomic displacement parameters (ADPs) and the placement of ordered solvent molecules, both of which are known to significantly impact model quality[3].

For ADP refinement in the guided search approach, we adopted a straightforward strategy: the ADPs from the PHENIX-refined model were copied and kept unchanged throughout the search. This ensures a fair comparison of *R*_free_ metrics between the methods. In the ensemble refinement approach, all ADPs were uniformly set to a small value of 5.0 Å^2^, encouraging conformational exploration without biasing the refinement toward any particular set of displacement parameters. While gradient-based optimization of B-factors is straightforward to implement using SFCalculator, preliminary trials showed that refining ADPs led to negligible improvements in the final metrics. This suggests that further investigation and more comprehensive testing are required to better understand and optimize ADP refinement within this framework.

### A better differentiable solvent model

The performed benchmarks revealed differences between the differentiable solvent mask implemented in SFCalculator and the Probe-Shrink method, particularly in the boundary regions. The Probe-Shrink method employs a non-differentiable “shrink” step which appears to better account for the hydration layer surrounding the macro-molecule. This adjustment is essential, as solvent molecules within the hydration layer are often ordered and cannot be adequately modeled using a flat solvent approximation. As a result, SFCalculator yielded slightly higher *R*_free_ values compared to PHENIX in scenarios where solvent contributions were prominent, especially at lower resolution. Importantly, as well, our comparisons with PHENIX were based on models without explicitly modeled solvent molecules because the generative models used here do not generate ordered solvent molecules.

These observations emphasize the necessity for a more advanced differentiable solvent model. Potential avenues for improvement include the development of differentiable morphological operations or the integration of a data-driven pre-trained masking approach, both of which could leverage the inherent differentiability of SFCalculator framework to enhance accuracy and performance.

The placement of ordered solvent molecules presents a further challenge, particularly when MD-based generative models, due to the stringent requirements of molecular topology. However, the differentiable nature of SFCalculator offers a promising avenue for addressing this issue as well. It could enable the development of a solvent placement model conditioned on both the structural model and experimental observables. Such a predictive approach would represent a desirable solution to the problem, leveraging gradient information to integrate solvent modeling seamlessly into the refinement process.

### Sensitivity to training scheme and mode collapse

The training process for current normalizing flow models can be particularly challenging, especially when incorporating objectives which involve the reverse Kullback-Leibler (KL) divergence. While forward KL-divergence encourages the learned distribution to be mass-covering, reverse KL-divergence drives the model to be mode-seeking. This tendency towards mode-seeking often results in mode collapse, where the learned distribution focuses on a narrow subset of the target distribution, neglecting other regions. To mitigate this issue, we used careful scheduling of the weights between these two objectives to balance mass-covering and mode-seeking behaviors, complicating the training process and limiting scalability. We emphasize that SFCalculator can be combined with any differentiable generative model and anticipate that these limitations will be overcome by the next generation of generative models.

## 4 Methods

### Structure factor from protein molecules, F_protein_

We adopt the direct summation method for the macromolecular part contribution. As the Fourier transform is linear, the overall contribution from the whole molecule equals to the summation of all atoms[57]:

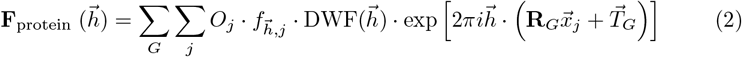

where *G* is the index of symmetry operations (appearing as rotation matrix 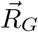 and translation vector 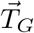 given the space group; *j* is the atom index, *O*_*j*_ is occupancy and 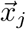 is the fractional coordinates of atom 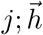 is the Miller index, 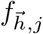 is the atomic scattering factor for the atom type of atom *j*, and DWF 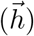 is the Debye-Waller factor.

The atomic scattering functions are approximated with the nine-parameter Gaussian summation[12], which is used in nearly all protein crystallography programs to compute the wavelength-independent atomic scattering factor (or atomic form factor) *f*_**S**_ as a function of the scattering angle *θ* :

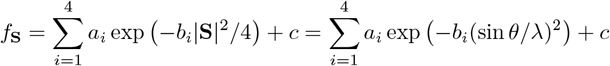

Here, *a*_*i*_, *b*_*i*_, and *c* are the Cromer-Mann coefficients. These coefficients are tabulated in the International Tables for Crystallography[53] In SFCalculator, atomic structure factor coefficients are accessed through GEMMI[67]. Additionally, to handle cryo-EM datasets, SFCalculator supports a “cryoem” mode in which the atomic structure factors are calculated using electron scattering factors parameterized as five Gaussians[53].

The Debye-Waller factor accounts for atomic displacement caused by thermal vibrations or structural disorder which results in slight positional variations of atoms within each unit cell, leading to additional phase differences. In crystallography, this effect is commonly parameterized as the B-factor. In isotropic case, the DWF becomes a standard Gaussian B-factor exponential:

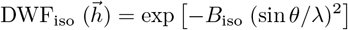

In the anisotropic case, the displacement is usually parameterized with a symmetric matrix *U*_*w*_, transforming the DWF into:

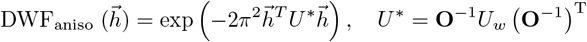

where **O**^−1^ is the deorthogoniazation matrix of the unit cell. SFCalculator is able to handle both isotropic and anisotropic parametrizations.

### Differentiable bulk solvent contribution F_solvent_

Calculating the solvent mask in a differentiable manner is a non-trivial task. The widely used probe-shrink method generates the solvent mask in a semi-localized way[28], utilizing van der Waals atomic radii *r* along with two parameters, *r*_*probe*_ and *r*_*shrink*_. While this approach achieves good agreement with experimental data, it involves non-differentiable operations such as rounding for discretization. Here, we propose an algorithm to approximate the solvent mask in a differentiable manner, as illustrated in Fig. 2a.

#### Algorithm 1

Differentiable solvent mask approximation

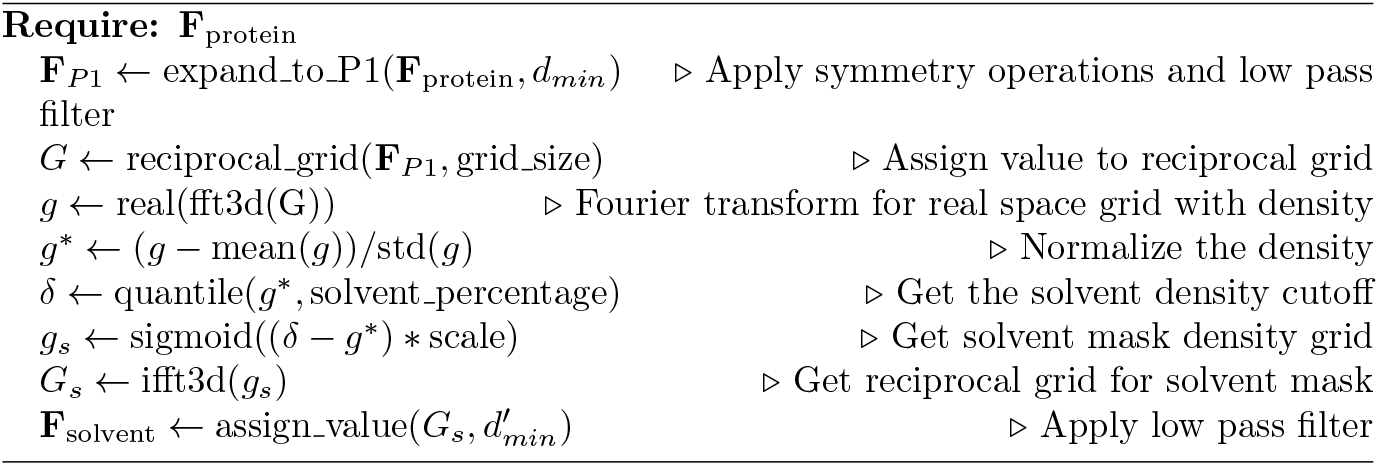

The complex protein structure **F**_protein_, calculated as described earlier, is first subjected to a Fourier transform to generate a protein density map. Subsequently, a sigmoid operation is applied after subtracting the density cutoff to produce a binary-like solvent mask map. This map is then processed through an inverse Fourier transform to calculate the structure factors contributed by the bulk solvent. It is important to note that two low-pass filters are applied in conjunction with the Fourier and inverse Fourier transforms to suppress high-frequency noise in the solvent mask map. Resolution cutoff 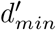 in the inverse Fourier transform is set as 3Å to be consistent with PHENIX default. This operation is conceptually aligned with the solvent flatness constraints previously proposed[59].

The algorithm incorporates three key hyperparameters: the low-pass filter cutoff *d*_*min*_, the solvent percentage, and the sigmoid scale. The methodology for determining these hyperparameters and the impact of their adjustments on the performance of SFCalculator are included in Appendix B.2.

### Scaling

After obtaining the two contributing components, **F**_protein_ and **F**_solvent_, additional parameters are introduced to scale the bulk-solvent model and align it with the experimental observables for accurate target function calculation. The scaling parameters in SFCalculator are initialized following the strategies outlined in [2], with enhancements achieved through gradient descent-based optimization.

The total model structure factor is defined as:

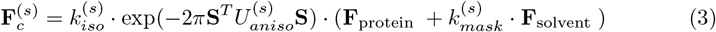

where the 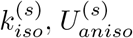 and 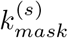 are scaling parameters in each resolution bin. The purpose of binning is to group data with common features, enabling each group to be characterized by a shared set of parameters. In this context, the primary parameter is the resolution *d* of reflections. SFCalculator employs a binning scheme that divides the resolution range uniformly on a logarithmic scale, ln(*d*) [65]. This approach ensures that higher-resolution bins contain more reflections than lower-resolution bins, while allowing for finer binning at low resolution without increasing the total number of bins. As highlighted in [2], the dependence of scale factors on resolution is approximately exponential. By using logarithmic binning, the variation of scale factors between bins is rendered more uniform, enhancing the algorithm’s effectiveness.

The scaling parameters are ideally determined by minimizing the following least-square residues:

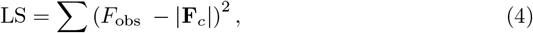

To achieve fast and reliable convergence, we employ the root-finding and linear system-solving methods proposed in [2] to initialize the scaling parameters. Details of these algorithms are provided in the Appendix B.3. Following this initialization, we leverage the autodifferentiation back-end of our SFCalculator implementation to perform a few steps of gradient descent-based optimization, using either the ADAM optimizer[32] or L-BFGS[37]. This process minimizes the target function (Equation 4) to compute the final scaling parameters.

### Molecular Replacement

The goal of molecular replacement is to determine the optimal rotation and translation of a molecule to best align with the experimental observables. Performing a full six-dimensional search, however, is computationally prohibitive. By decomposing the rotation and translation searches, the number of evaluations can be significantly reduced. Hierarchical grid search[68, 69] allows for additional efficiency gains.

Our molecular replacement method begins with a rapid packing score (see Appendix B.1) search performed on a coarse grid of center-of-mass (COM) positions within the asymmetric unit, using a random initial pose. This step identifies a candidate COM with the lowest clash score. The selected COM, in combination with the initial random rotation, is then used to construct an initial model and determine the scale factors. The identified COM is subsequently carried forward to the rotation search phase. It is important to note that the following hierarchical grid search remains indispensable despite the availability of a fully differentiable backend capable of gradient-based optimization. This necessity arises from the highly rugged landscape of the target function, which limits the radius of convergence for gradient descent. Therefore, identifying a suboptimal candidate through the hierarchical search is a critical prerequisite. The next phase involves a rotation search on a hierarchical SO(3) grid, leveraging Patterson functions to determine a suboptimal rotation matrix. This matrix is then utilized in a hierarchical translation search, which identifies a suboptimal translation vector. The pipeline culminates in a gradient descent optimization step, refining the pose to achieve the final solution.

### Hierarchical rotation search

The decomposition of the rotation and translation search is achieved using Patterson functions [56]. The Patterson function is defined as the autocorrelation of the electron density, effectively representing a map of interatomic distance vectors:

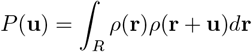

Leveraging the convolution theorem, the Patterson map can be expressed as the Fourier transform of the squared magnitudes of the structure factors, which are derived directly from experimental data:

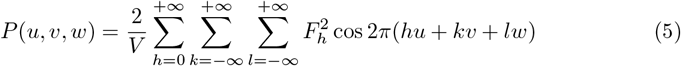

#### Algorithm 2

Hierarchical Rotation Search

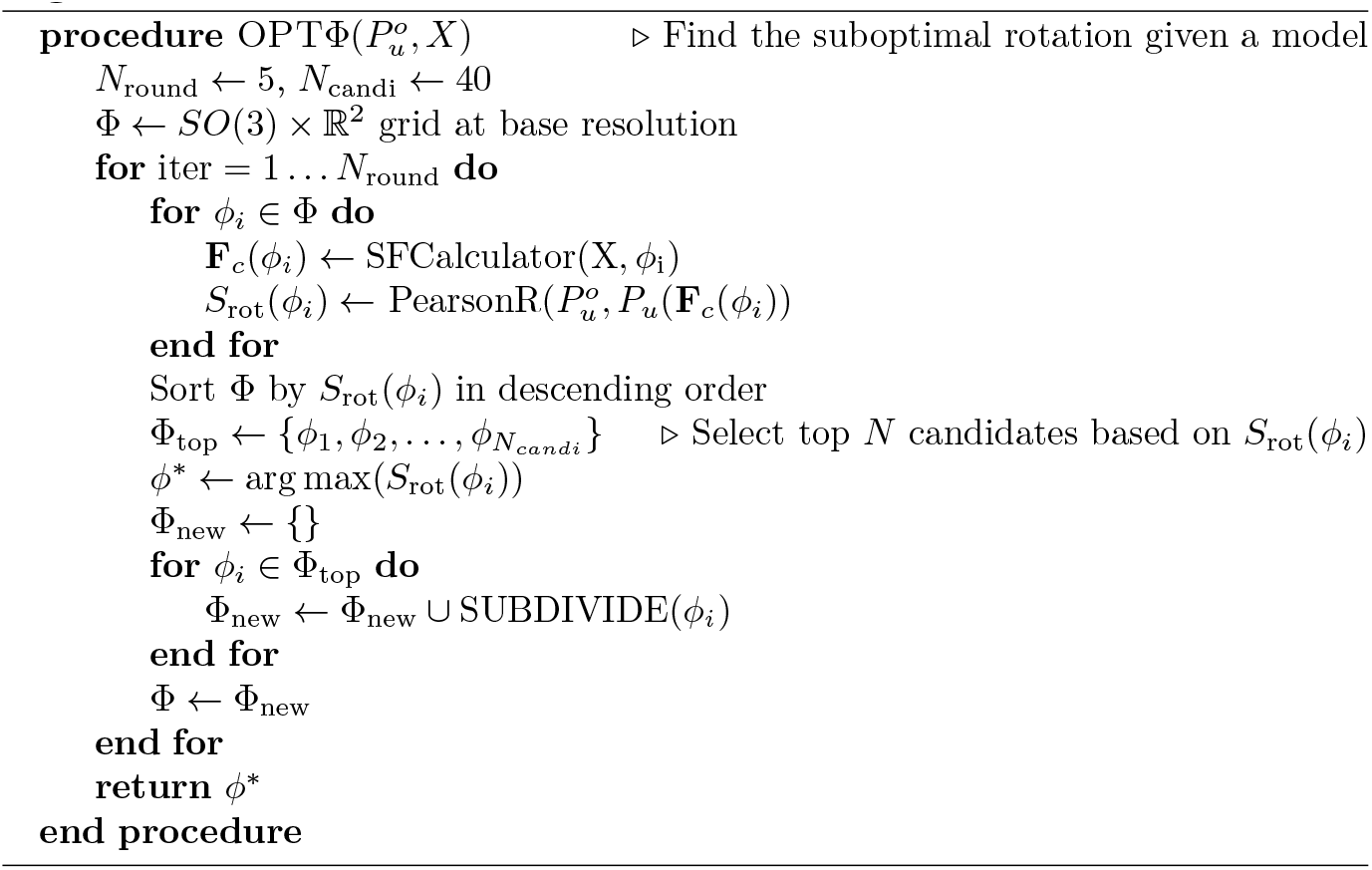

As the autocorrelation of the electron density, the Patterson map inherently captures the interatomic features of the system. Specifically, by focusing on an appropriate range of Patterson vectors that primarily represent intramolecular pairwise distance features, the map becomes relatively insensitive to translational changes. Consequently, using the Patterson function as the target allows for the decomposition of the rotational and translational searches, significantly simplifying the molecular replacement process.

In practice, we use the following correlation function as the scoring function for hierarchical rotation search:

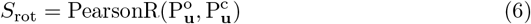

where 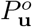 is the Patterson function calculated using experimental structure factor amplitudes [57], while 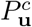 is the Patterson function calculated using the structure factor amplitudes of candidate models.

For the hierarchical grid search, we adopted the uniform multiresolution grids on SO(3) as described in[68], utilizing the Healpix [21] grid for the sphere and the Hopf fibration to uniformly lift the grid to *SO*(3). This approach is consistent with the method used in cryoDRGN[69]. The base grid on SO(3) consists of 576 orientations, with a spacing of 30^°^. In each iteration, the 40 grids with the highest *S*_rot_-scores are selected and further subdivided into higher-resolution neighboring grids, with each grid generating 8 neighbors. Typically, five rounds of refinement are performed, culminating in a final orientation resolution of 0.92 degrees. This hierarchical approach requires a total of 1,856 evaluations, significantly fewer than the brute-force search involving 36,864 rotations at a spacing of 7.5^°^, while achieving a much finer rotation spacing.

### Hierarchical translation search

The concept of the hierarchical translation search mirrors that of the hierarchical rotation search, with the primary difference being that the subdivision of grids is performed in fractional coordinates using interpolation. At each level, each grid point generates 3^3^ − 1 = 26 neighboring grids in the next level. Additionally, since a suboptimal rotation matrix has already been determined from the preceding search, there is no need to rely on a Patterson function-based scoring method. Instead, we utilize the correlation of structure factor magnitudes, which reduces the computational cost by avoiding the calculation of the Patterson function. The correlation is quantified using the Pearson correlation coefficient between observed structure factor amplitudes, *F*_*obs*_, and structure factor amplitudes calculated from the model, *F*_*c*_:

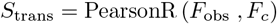

The translation search can still be computationally expensive, even with a hierarchical strategy, due to the size of the unit cell. However, one advantageous property of crystallographic symmetry can significantly reduce the computational effort: the polar axis. The polar axis, present in specific space groups, is an axis along which translation operations do not affect the structure factor magnitudes. This means that any point along the polar axis can be arbitrarily chosen as the origin, eliminating the need to perform a search along that axis. SFCalculator implementation fully supports this functionality, allowing for more efficient translation searches.

### Gradient-Descent Rigid Body Refinement

Once the suboptimal rotation *ϕ*_0_ and translation *υ*_0_ have been determined, we assume that they lie within the radius of convergence for gradient descent. Gradient-based optimization is then performed using the following target function:

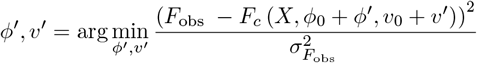

The optimal pose is subsequently defined as:

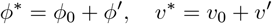

For this optimization, rotations are parameterized as quaternions to facilitate unconstrained optimization.

### Crystallographic *R* factors

*R* factors are statistical metrics used in crystallography to assess the quality of a structural model, calculated by comparing the model-predicted structure factors to crystallographic data, with lower values indicating better agreement.

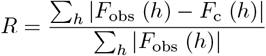

Where *h* indicates triplets of so-called Miller indices that indicate the spatial frequencies of the corresponding Fourier components. Typically, two separate *R* factors are calculated during structure refinement: *R*_*work*_ is calculated over Miller indices included in refinement, while *R*_*free*_ is a cross-validation metric calculated using a reserved subset of experimental data not used during refinement. A smaller gap between *R*_*work*_ and *R*_*free*_ indicates less overfitting.

### Training for physical prior embedded model

The physical prior is represented by the Boltzmann distribution, *p*(*x*) = *e*^−*E*(*x*)/*RT*^ /*Z* at temperature *T*, with *R* the universal gas constant; *Z* is the partition function. To approximate the target distribution using our normalizing flow model, we utilize two training objectives:

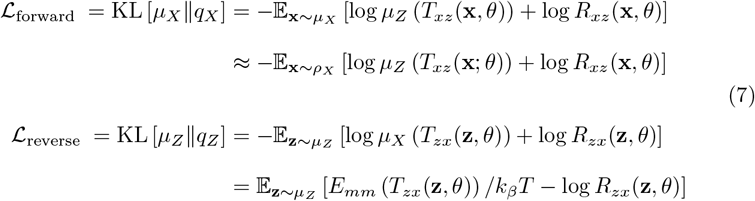

Here:

1. ℒ_forward_ corresponds to the negative log-likelihood of the training dataset, as in most generative models. Training samples are typically prepared by running molecular dynamics simulations for 100 ns using an implicit solvent model. *µ*_*Z*_ represents the prior density distribution in the latent space, which is Gaussian in this case; *µ*_X_ represents the ground truth distribution of protein conformations, and we approximate it with *ρ*_*X*_, the observed density of MD samples.
2. ℒ_reverse_ is the energy-based term specific to cases where the (normalized) density function of the target distribution is known. It ensures that sampled conformations from the latent space exhibit physically reasonable energies. *E*_*mm*_ represents the molecular mechanics energy. It is important to note that most molecular mechanics force fields can be easily wrapped as differentiable energy functions, since the (negative) gradient of the energy with respect to the coordinates corresponds directly to the force. In the work, we employed OpenMM[16].

The training process begins by optimizing ℒ_forward_ alone to initialize the network. At this stage, generated samples resemble those in the molecular dynamics trajectories but often contain steric clashes or other nonphysical details, leading to high energies. Subsequently, the weight of ℒ_reverse_ is gradually increased to regularize the latent space, incorporating physical energy constraints and ensuring sampled conformations align with the Boltzmann-distributed target.

### Experimental data likelihood

The differentiability of SFCalculator enables a differentiable calculation of model structure factors **F**_*c*_, thus the likelihood target. We employ the following two objective likelihood functions:

1. Least Squares Loss:

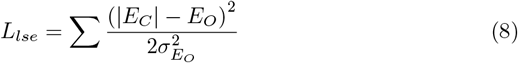
2. Negative Log-Likelihood from Rice distribution:

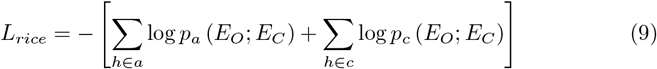

where, according to [54], the likelihoods *p*_*a*_ (*E*_*O*_; *E*_*C*_) for acentric reflections and *p*_*c*_ (*E*_*O*_; *E*_*C*_) for centric reflections are defined as:

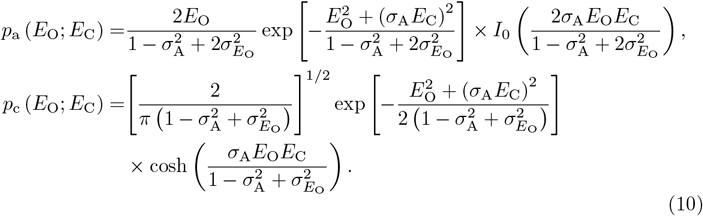

Here, *E*_*O*_, *E*_*C*_, and 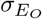 represent the normalized *F*_*o*_, *F*_*c*_ (such that ⟨*E*^2^ ⟩= 1), and *σ*_*F*_, respectively. *σ*_*A*_ describes the correlation between model and data, as defined in [54, 63].

During the guided search approach, gradient descent is performed with respect to a linear combination of *L*_*lse*_ and *L*_*rice*_. The weights of this combination are chosen empirically and depend on the specific system under consideration.

### Training for posterior approximation

The training objective for the normalizing flow model approximating the posterior is to minimize the following reverse Kullback–Leibler divergence:

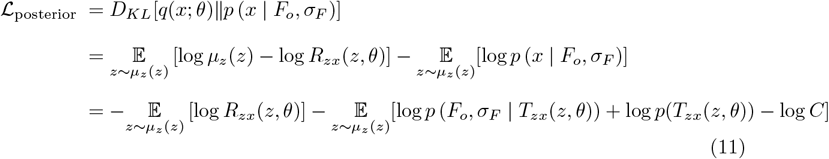

which is equivalent to maximizing the Evidence Lower Bound (ELBO) as defined in [55]. log C is the normalization constant.

By using the Boltzmann distribution as the physical prior, the energy term is incorporated, with the normalizing constant (partition function) absorbed into the constant term:

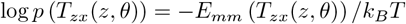

Thus, the objective ℒ_posterior_ is equivalent to ℒ_reverse_ as defined in Equation 7, with the addition of the negative experimental log-likelihood term − log *p* (*F*_*o*_, *σ*_*F*_ | *T*_*zx*_(*z, θ*)).

The training process begins by optimizing ℒ_forward_ alone to initialize the network. Once the initial training is complete, the weight of ℒ_posterior_ is gradually increased to regularize the latent space. This step incorporates both physical energy and experimental likelihood constraints, ensuring that the sampled conformations align with the posterior target.

In greater detail, during the training phase involving ℒ_posterior_, the negative log-likelihood term initially uses the least squares term *L*_*lse*_ as defined in Equation 8. Over time, this is gradually transitioned to *L*_*rice*_, as defined in Equation 9. This approach is adopted because, during the early stages of ℒ_posterior_ training, the energy term is often very large due to the presence of non-physical details in the conformations. Under these conditions, the least squares term *L*_*lse*_ is more suitable for matching the scale of the other terms in the loss function. Transitioning to *L*_*rice*_ later in the process ensures a more accurate incorporation of the experimental likelihood once the system has been sufficiently regularized.

## Code availability

SFCalculator has been implemented with PyTorch[52], JAX[8], and TensorFlow[1] backends to accommodate a broader range of users. The codebase is open-sourced and available at: https://github.com/Hekstra-Lab/SFcalculator.

Code related to molecular replacement, normalizing flow construction, training and refinement is maintained in a separate repository, `deeprefine`, which can be accessed at: https://github.com/minhuanli/deeprefine.

## Acknowledgements

We thank Tom Terwilliger, Randy Read, Alisia Fadini, members of the Hekstra Lab and the Molecular Biophysics and Integrated Bioimaging at Lawrence Berkeley National Lab for fruitful discussions. This work was supported by a graduate fellowship of the Eric and Wendy Schmidt Center to M.H.L., a Career Award at the Scientific Interface from the Burroughs Wellcome Fund to K.M.D., and the NIH Director’s New Innovator Award (DP2-GM141000) to D.R.H. Computations were run on the FASRC Cannon cluster supported by the FAS Division of Science Research Computing Group at Harvard University. We acknowledge the open source software projects used in this work, including reciprocalspaceship[22], GEMMI[67], and healpy[70].

## Declarations

The authors declare no competing interests.

## Author Contributions

M.H.L., K.M.D. and D.R.H. conceived the project. M.H.L. developed all code. M.H.L. and D.R.H. performed the analysis and drafted the manuscript. All authors edited the paper.

## Appendix A Supplementary Figures

**Fig. A1.**
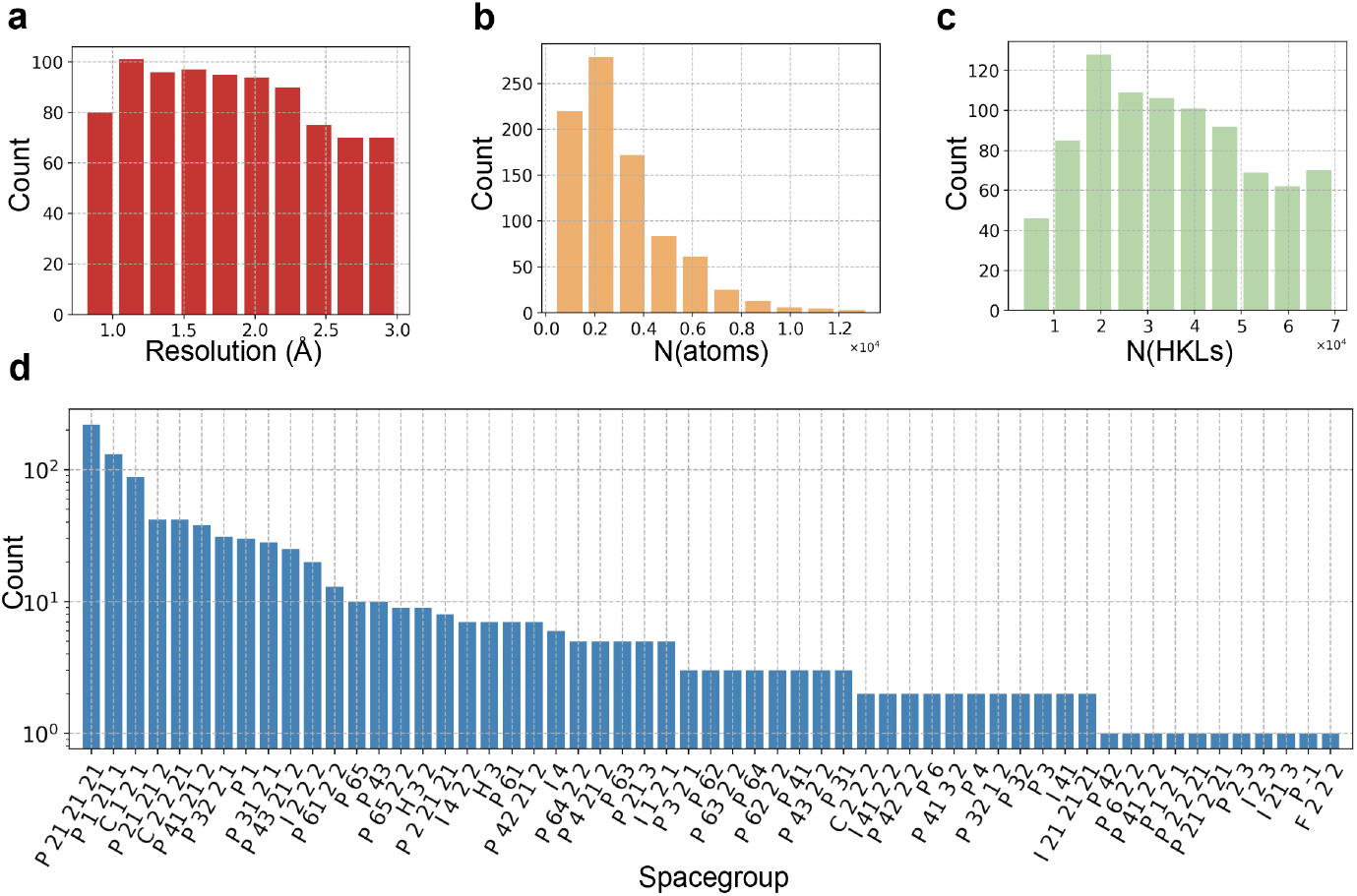
Statistics of PDB entries used as benchmark datasets. Benchmarking was conducted across 868 PDB entries, encompassing data resolutions ranging from 0.8 Å to 3.0 Å (**a**), diverse molecular size (**b**), various number of reflections (**c**), and various space groups (**d**).

**Fig. A2.**
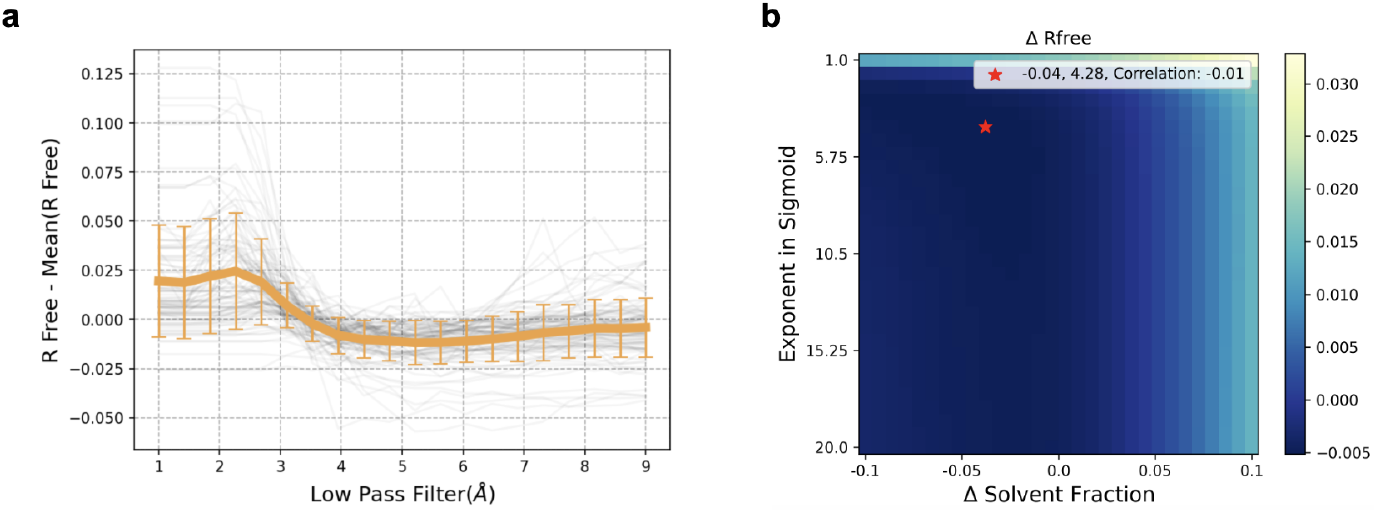
**Sweep of hyperparameters used in the differentiable solvent mask**, with results based on statistics from 300 PDB entries. The final *R*_free_ value is used as the primary evaluation metric, with lower values indicating better performance. **a**. Effect of the low-pass filter cutoff. The optimal range is found to be 4-6 Å, with 5 Å selected as the default for its balance of accuracy and generality. **b**. The normalized voxel density distribution is passed through a sharp sigmoid function with inflection point dependent on the estimated solvent fraction (see Algorithm 1). Here we swept an offset, Δ between the sigmoid midpoint and estimated solvent fraction as well as the exponent of the sigmoid function. The results demonstrate that the solvent percentage algorithm is robust, with minimal sensitivity to changes as Δ within the range of −5% to +2%. Similarly, the exponent of the sigmoid function is not important as long as it remains sufficiently large. A higher exponent ensures a more binary solvent mask. The default value of 10 was chosen for its consistency across tests.

**Fig. A3.**
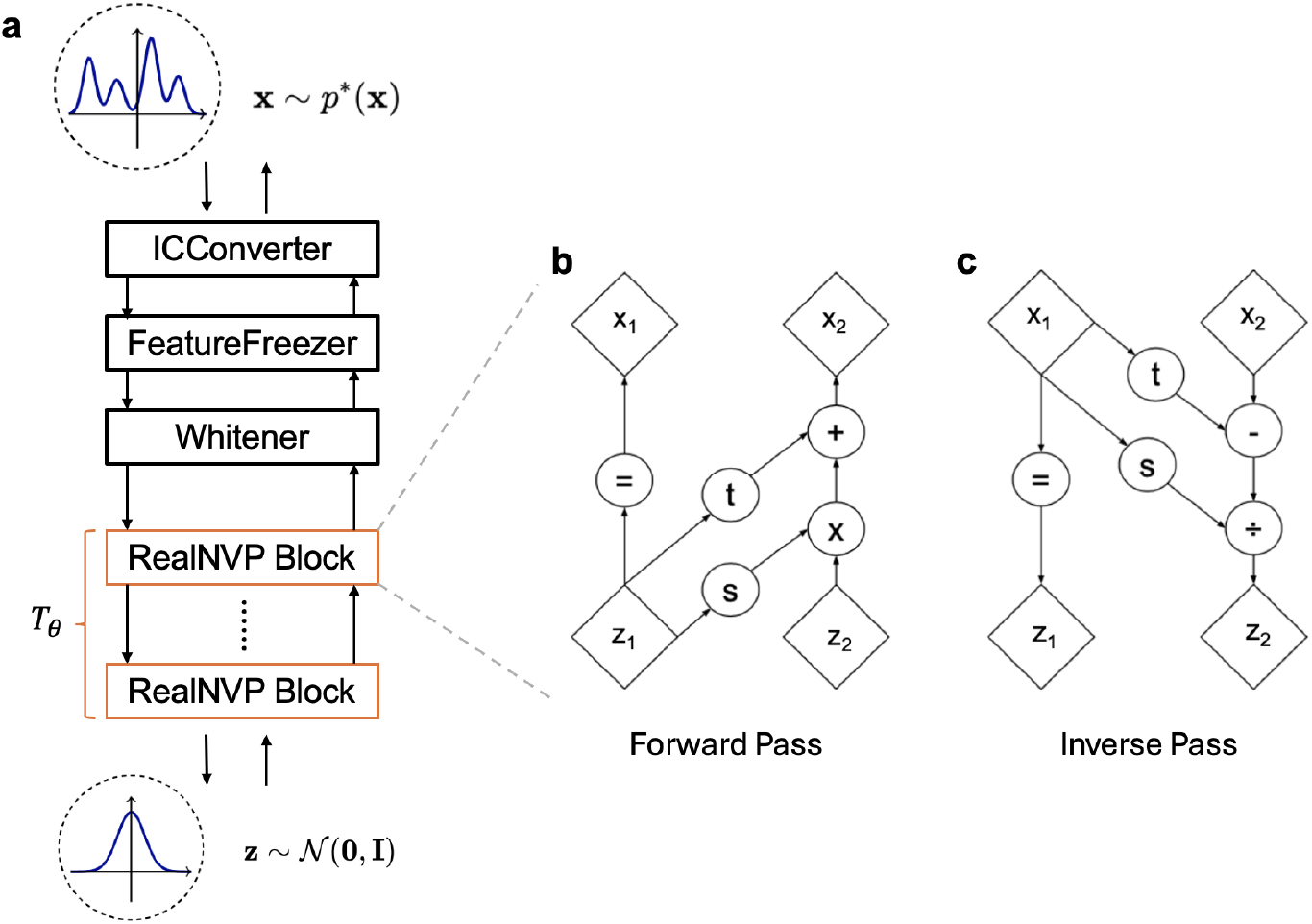
Architecture of the normalizing flow model. **a**. Schematic representation of the model pipeline (see section B.4 for implementation details). Cartesian coordinates are first converted into internal coordinates using the ICConverter, followed by the FeatureFreezer, which locks features with low variability, such as bond lengths. Next, the Whitener normalizes the features and removes the six degrees of freedom associated with global rigid-body transformations. Finally, the data is processed through a stack of trainable RealNVP blocks. **b**. Each RealNVP block consists of two coupling layers that apply invertible transformation with channels swapped, combining scaling and the addition of a constant offset to one subset of the input vector (*z*_2_), conditioned on the remaining subset (*z*_1_). **c**. The inverse transformation of the coupling layer, illustrating the reversibility of the model. Panel **b** and **c** are adapted from reference[15].

**Fig. A4.**
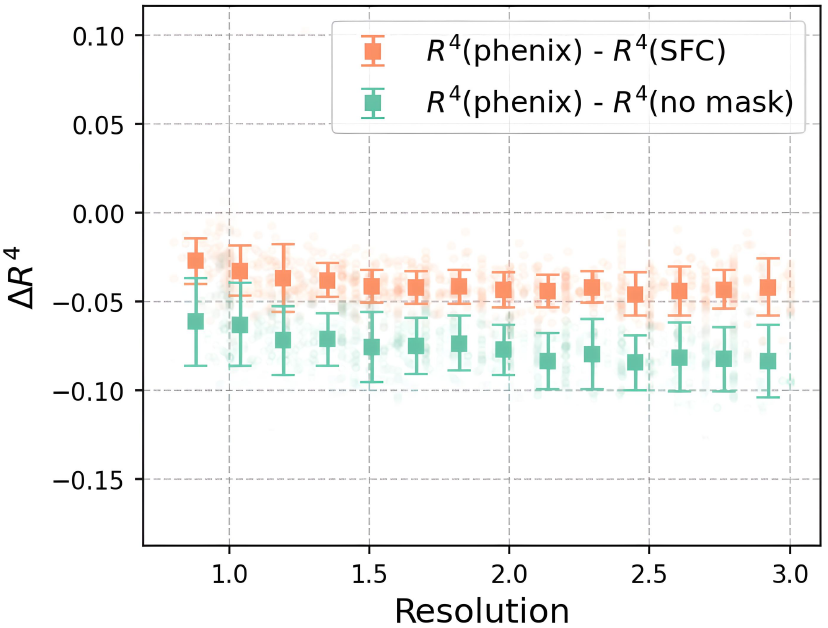
Statistics of *R*^4^ comparisons between SFCalculator, PHENIX, and the case with no solvent mask applied. *R* factors were calculated over reflections at resolutions ≥ 4 Å comparing observed structure factors against ones calculated from the 868 corresponding deposited models using SFCalculator or PHENIX with solvent mask, or using just the protein structure factors. Results were binned across datasets by their resolution, with error bars indicating standard deviations. The results clearly demonstrate that the differentiable solvent mask used in SFCalculator outperforms the absence of a mask across all ranges. However, due to its handling of the hydration layer, the Probe-Shrink method, the default in PHENIX, achieves better accuracy than SFCalculator.

## Appendix B Detailed methodologies

### B.1 Fast voxel rendering for solvent fraction calculations

The concept of fast solvent percentage calculation involves estimating the occupancy of protein molecules on a coarse unit cell grid, typically with a spacing of *z*= 4.5 Å. For this purpose, we adopt the occupancy function proposed in PyUUL[48]:

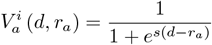

Here, 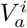 represents the fraction of volume *i* occupied by an atom *a* with radius *r*_*a*_ at a distance *d* from its center. The parameter *s* controls the steepness of the decay. The total volume occupancy for each grid point is calculated as:

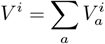

If *V*^*i*^ > *c*, where *c* is the cutoff fraction, the volume is classified as “occupied.” The solvent percentage is then computed as:

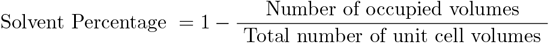

For the steepness parameter *s*, we adopt an empirical formula:

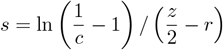

This ensures that any atom center within *z*/2 of the volume center will occupy the volume. By default, we set *c* = 10^−3^, resulting in a large *s* value to ensure rapid density decay. This ensures that a volume is considered occupied only when it is significantly filled by an atom, rather than being the result of a cumulative effect from multiple atoms contributing low-density values.

The same algorithm is employed for calculating packing scores and clash scores in the molecular replacement pipeline. The packing score is defined as the fraction of occupied volumes, while the clash score is defined as the fraction of volumes occupied by two or more atoms.

### B.2 Hyperparameters in solvent mask calculation

To evaluate the impact of hyperparameters in the solvent model, we conducted a systematic sweep using 300 PDB entries, with the final *R*_free_ value serving as the primary evaluation metric (lower values indicate better performance), as shown in Fig. A2. The analysis of the low-pass filter cutoff revealed that the optimal range is between 4 and 6 Å, with 5 Å selected as the default for its balance between accuracy and generality. Additionally, we examined the effects of (1) applying an offset to the solvent fraction estimate calculated above for use in the solvent mask calculation (see Algorithm 1) and (2) varying the exponent in the sigmoid function. The solvent fraction algorithm proved robust, exhibiting minimal sensitivity to deviations in the solvent percentage offset within the range of −5% to +2%. Similarly, the exponent in the sigmoid function was found to be effective as long as it remained sufficiently large to ensure a more binary solvent mask. A default exponent value of 10 was chosen based on its consistency across the tests.

### B.3 Robust initialization of scaling parameters

We initialize the *k*_*iso*_, *k*_*mask*_ and *U*_*aniso*_ in a fast and robust way following the reference[2], with necessary modifications.

First, we initialize *k*_*mask*_ and *k*_*iso*_ by solving the following overdetermined minimization problem for each resolution bin *s*:

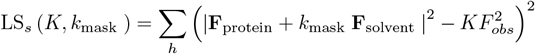

where 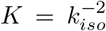, and in each resolution bin *k*_*iso*_ and *k*_*mask*_ are fixed for each reflection *h*. Minimizing the above equation lead to initializing *k*_*mask*_ by root finding of the following equation:

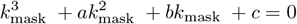

where:

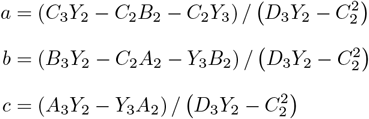

and:

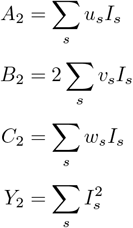

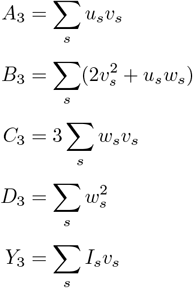

where *w* = |**F**_solvent_ |^2^, *v* = (**F**_protein_, **F**_solvent_), *u* = |**F**_protein_|^2^, and 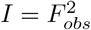

If no positive root exists, *k*_mask_ is assigned a zero value, which implies the absence of a bulk-solvent contribution. If several roots with *k*_mask_ ≥ 0 exist then the one that gives the smallest value of LS_*s*_ (*K, k*_mask_) is selected. Once *k*_*mask*_ is determined, *k*_*iso*_ is calculated with:

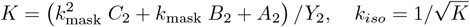

Once we have initialized *k*_*mask*_ and *k*_*iso*_, the remaining *U*_*aniso*_ is determined by solving a linear system:

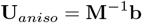

where **M** = Σ_s_ **V** ⊗ **V, V** = (*h*^2^, *k*^2^, *l*^2^, 2*hk*, 2*hl*, 2*kl*)^*t*^, **b** = − Σ_s_ *Z***V**, and *Z* = [1/ (2*π*^2^)ln *F*_obs_ (*k*_*iso*_ |**F**_protein_ + *k*_solvent_ **F**_mask_ |)^−1^]

### B.4 Architecture of the Normalizing Flow model

The schematic architecture, illustrated in Fig. A3a, is composed of several key blocks, which we describe in detail below:

#### ICConverter

We adopt the mixed coordinate transformation layer described in [47]. In this approach, the Cartesian coordinates of macromolecules are partitioned into backbone atoms (N, *C*_*α*_, C for each residue) and auxiliary atoms. The backbone atoms are retained as Cartesian coordinates (*x*_*C*_), while the auxiliary atoms are converted into internal coordinates (*x*_*I*_) as follows: for each auxiliary atom *i*, three “parent” particles *j, k*, and *l* are defined. The Cartesian coordinates of particles *i, j, k*, and *l* are then transformed into distance, angle, and dihedral coordinates (*d*_*ij*_, *α*_*ijk*_, *ϕ*_*ijkl*_). Consequently, the Cartesian coordinates *x* are transformed into a combination of Cartesian and internal coordinates, expressed as *x* → [*x*_*C*_, *x*_*I*_].

The inverse transformation is straightforward. Auxiliary atoms are positioned sequentially: first, those whose parent particles are entirely within the Cartesian set, followed by particles whose parents have just been placed, and so on. The conversion from internal coordinates to Cartesian coordinates is performed using the NeRF algorithm [51].

#### FeatureFreezer

The FeatureFreezer module is newly introduced to lock features with low variability, thereby reducing the dimensionality of the problem. This is achieved through the following steps:

1. Identify the features to be frozen based on statistical analysis of the training set.
2. Exclude these features during the forward pass.
3. Reintroduce them in the inverse pass using their statistical mean values.

The selection of frozen features is governed by the following rules:

1. Cartesian signals are never frozen.
2. Distance signals (covalent bonds’ lengths) are frozen if the ratio of standard deviation to the mean (std/mean) is less than 0.05.
3. Angle signals, represented as **v**_*i*_ = [sin(*θ*), cos(*θ*)], are frozen if the circular concentration coefficient *c* = |Σ**v**_*i*_|/Σ|**v**_*i*_| exceeds 0.996, corresponding to a standard deviation of approximately 5 degrees.

#### Whitener

The Whitener layer employs Principal Component Analysis (PCA) to normalize feature scales and remove their means. Two separate PCA operations are applied:

1. For Cartesian signals (*x*_*C*_), PCA performs whitening and removes the six degrees of freedom associated with global rigid-body transformations.
2. For internal coordinates, PCA is applied independently to normalize these features.

The whitening and inverse whitening matrices are stored, enabling efficient application during both the forward and inverse passes.

#### RealNVP block

The RealNVP block incorporates trainable parameters to define and shape the transformation. Each RealNVP block consists of two stacked coupling layers, with channels swapped between the layers to ensure that both channels are transformed [15, 47]. A schematic representation of a single coupling layer is shown in Fig. A3b and c. The forward transformation is defined as:

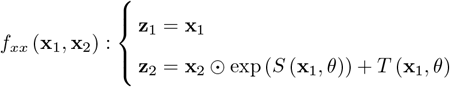

where ⊙ is element-wise multiplication, and the log-determinant of the Jacobian given by:

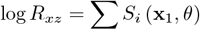

The inverse transformation operates as follows:

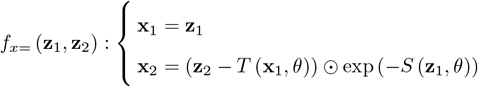

with the corresponding log-determinant:

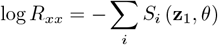

This setup ensures that the transformation remains invertible while allowing fast computation of the log-determinant for both forward and inverse passes.

